# Prolonged anesthesia alters brain synaptic architecture

**DOI:** 10.1101/862334

**Authors:** Michael Wenzel, Alexander Leunig, Shuting Han, Darcy S. Peterka, Rafael Yuste

## Abstract

Prolonged medically-induced coma (pMIC), a procedure performed in millions of patients worldwide, leads to cognitive impairment, yet the underlying brain mechanism remains unknown. No experimental studies of medically-induced coma (MIC) exceeding ~6 hours exist. For MIC of less than 6 hours, studies in developing rodents have documented transient changes of cortical synapse formation. However, in adulthood, cortical synapses are thought to become stabilized. Here, we establish pMIC (up to 24 hrs) in adolescent and mature mice, and combine repeated behavioral object recognition assessments with longitudinal two-photon imaging of cortical synapses. We find that pMIC affects cognitive function, and is associated with enhanced synaptic turnover, generated by enhanced synapse formation during pMIC, while the post-anesthetic period is dominated by synaptic loss. These results carry profound implications for intensive medical care, as they point out at significant structural side effects of pMIC on cortical brain synaptic architecture across age levels.

---

In humans, cognitive impairment due to prolonged medically induced coma (pMIC) represents an enormous clinical and socio-economic burden affecting millions of patients worldwide^1–5^. In adulthood, clinical trials have proven that intensive care unit (ICU) survivors frequently suffer from lasting cognitive impairment^1,6^, yet its pathophysiology and neuromorphopathological underpinnings have remained elusive. In early life (childhood and early adolescence), when the brain is highly plastic, MIC (max. 6 hrs tested in rodents to date) has been shown to result in synaptic changes and long-term cognitive impairment^7–9^. However, basic animal research suggests that during later adolescence and adulthood, dendrites and dendritic spines become stabilized under physiological conditions^10–12^, and short-term MIC^13^. Whether this notion holds true for pMIC, is unknown.

There has been a paucity of experimental studies of structural and functional consequences of pMIC, so our first objective was to create a robust platform for measurement in mice. We combined a mobile anesthesia setup (Fig. 1a), chronic two-photon imaging of cortical synapses^14^ (Fig. 1b) through a reinforced thin skull window^15^ (Fig. 1b), and repeated behavioral assessment^16^ (Fig. 1c) into an integrated experimental framework (Fig. 1d, please see methods). Two imaging sessions were carried out to determine baseline 3-day (‘d’) synaptic turnover (d1, d4). Immediately after d4 imaging, pMIC was initiated. To document synaptic changes across pMIC, a third imaging session was carried out immediately upon pMIC discontinuation (d5). Three days post pMIC, mice were imaged once again to determine post-anesthetic 3-day synaptic turnover (d8). Additionally, a novel object recognition test^16^ was performed during baseline conditions (d4), and repeated after recovery from pMIC (d7, fig. 1c).

**Figure 1.**
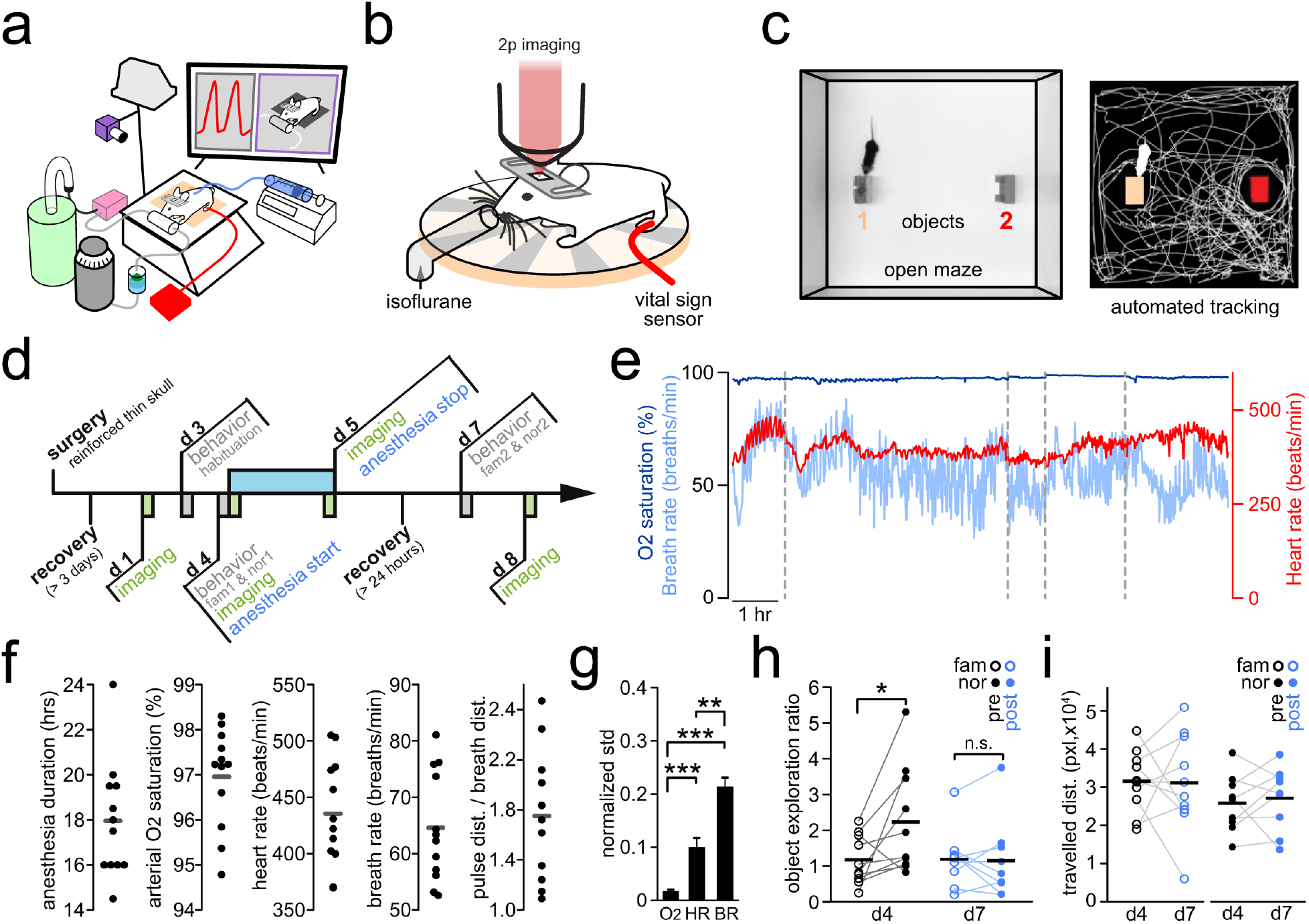
Experimental design and pMIC-induced behavioral deficits. **a)** Sketch of the anesthesia setup for pMIC. The mouse is placed on a warming pad on a tiltable platform with its nose resting in a custom nose piece. Isoflurane is evaporated (grey), and delivered in humidified air/oxygen. Waste gas is removed via a suction pump (magenta) and trapped in a charcoal filter (turquoise). Nutritional supply is delivered via a subcutaneous line by a programmable perfusion system (blue, right). In addition to the warming pad, an infrared lamp is placed above the setup. Vital parameters are measured using a thigh sensor (red). Vital parameters and clinical status are video recorded and live-streamed in a local network (camera above the setup, cmputer display). **b)** for individual imaging sessions (max. 30 min), the mouse is placed under a two-photon microscope, on a warming pad, under isoflurane anesthesia, while vital signs are being monitored using a thigh sensor. **c)** open maze for behavioral testing. A camera is mounted atop the maze. Videos are recorded during object familiarization and novel object recognition, and the mouse behavior is automatically tracked using custom written MATLAB code. **d)** Experimental workflow, fam = familiarization, nor = novel object recognition test. **e)** Basic vital parameter long-term monitoring of an anesthetized mouse, with intermittent breaks (dotted lines, e.g. for moving the sensor from one extremity to another). Faulty measurements for technical reasons (e.g. tissue compression) are excluded. Displayed recording length ~12 hrs. **f)** measured set of vital parameters, mean values (n = 12 mice), overall mean value displayed as horizontal grey line. dist. = distension. **g)** Comparison of the core vital parameter normalized mean standard deviations (n= 12 mice). Unpaired t-test with Welsh’s correction: O2 saturation vs. heart rate p=0.001, O2 saturation vs. breath rate p<0.0001, heart rate vs. breath rate p=0.0002. **h)** basic cognitive testing: relative object exploration time ratio (2 objects, 50% vs. 50%= 1) during familiarization (“fam”, circles, 2 identical objects), followed by novel object recognition (“nor”, filled circles, 1 of the identical objects replaced by a novel object) measured during baseline (d4), and post recovery from pMIC (d7). Paired t-test: fam (d4) vs. nor (d4) p=0.0451 (n=10 mice), fam (d7) vs. nor (d7) p=0.823 (n=9 mice). **i)** the reduction in “nor” performance can not be explained by reduced locomotion, travelled distances (in pixels) prior and post pMIC during fam (circles), and nor (filled circles), are consistent. Depiction of statistical significance in fig. 1: *p<0.05, **p<0.01, ***p<0.001. All bar plots show mean ± s.e.m..

We used the inhalatory anesthetic isoflurane due to its fast kinetics (order of minutes) and thus maximal adjustability of anesthetic depth^17^. We tried to avoid an FiO_2_ (fraction of inspired oxygen) >50% throughout pMIC, as high FiO_2_ values have been associated with increased mortality in human intensive care^18^. Vital parameters (peripheral O_2_ saturation, heart rate, breath rate, pulse distention, breath distention) were closely monitored using an infrared thigh sensor (Fig. 1e). When developing the experimental protocol, we successfully performed extended anesthesia in mice for up to 40 hours, yet for reasons of experimental feasibility (continuous presence of an experimenter required), we carried out pMIC on 12 adult mice (age: 7.7 ± 1.5 months s.e.m.) with a mean duration of 18 hours (17.96 ± 0.74 s.e.m.). Cardiorespiratory instability, or maximum pMIC duration of 24 hours were chosen as primary pMIC discontinuation criteria. Fluid and nutritional supply was ensured by administration of glucose solution (1%, in PBS) via a subcutaneous line at 0.1-0.2ml/hr (total volume 3.46 ± 0.35 ml s.e.m.). Under steady state isoflurane concentrations of 0.5-1.5% ppa, mice displayed continuously high peripheral oxygen saturation (96.96 ± 0.32 % s.e.m.), heart rates between 350-500 beats per minute (435 ± 14 s.e.m.), breath rates of 50-80 excursions per minute (65 ± 3 s.e.m.), and a pulse/breath distension ratio above 1 (1.75 ± 0.16 s.e.m.) (Fig. 1e-f). Sufficient oxygenation (>90%) was achieved with little variance across a wide range of heart and breath rates (Fig. 1e-g). Relative variance of heart and breath rate was much greater than O_2_ saturation. While heart and breath rate co-varied over time (Fig. 1e), breath rate showed the largest variance of all monitored vital parameters (Fig. 1g), and most accurately clinically reflected anesthetic depth. Upon pMIC discontinuation, mice usually rapidly displayed an acceleration of heart and breath rate, and soon reacted to tactile stimulation. We defined the period between isoflurane discontinuation and first appearance of the righting reflex as the wake-up time (9.65 ± 1.41 min s.e.m.). After waking up, all studied mice soon showed apparently normal movement and body coordination as well as typical grooming and feeding behavior. To evaluate whether pMIC results in basic behavioral changes, mice were repeatedly (prior and post pMIC) subjected to a novel object recognition test (‘NOR’, fig. 1c). In line with previous literature^16^, mice spent significantly more time exploring novel objects post familiarization during baseline conditions (Fig. 1h, left). However, this effect was abolished post recovery (>24hrs) from pMIC (Fig. 1h, right). Importantly, this effect could not be explained by a simple post-anesthetic reduction of navigation through the maze, as pre- and post-anesthetic total distances travelled during both familiarization and NOR were similar (Fig. 1i).

Next, we set out to study whether pMIC affects synaptic architecture. To this end, we carried out repeated structural two-photon imaging of YFP-labeled dendrites and synapses (up to ~150μm beneath the pial surface) of layer 5 pyramidal neurons^14^ in 6 of the 12 adult mice undergoing pMIC (Fig. 2a). We chose a minimally invasive approach, using a reinforced chronic cranial window over the thinned skull to minimize possible effects from invasive surgery. However, this reduces overall imaging fidelity, and we were not able to consistently distinguish between filopodia and dendritic spines across all imaging sessions in all mice. Therefore, we chose to focus on clearly visible synaptic protrusions, regardless of morphological classification. In 6 mice, we assessed a total of 1254 synaptic protrusions (209 ± 40 s.e.m.) on 238 dendritic compartments (40 ± 9 s.e.m.). We noticed that across all anesthetized mice and imaging sessions, dendrites remained stable (Fig. 2b-c). However, pMIC-associated synapse dynamics showed considerable alterations (Fig. 2d). In comparison to the 1-day total synapse turnover in adult controls (n=6, total synapse count: 1113 [185.5 ± 50 s.e.m.]), pMIC-associated synapse turnover was more than doubled (Fig. 2e). Both synaptic gain and loss were significantly enhanced during pMIC, with the gain of synapses outperforming their loss (Fig. 2f). When comparing pre- and post-anesthetic 3-day synaptic turnover, turnover remained significantly enhanced after pMIC (Fig. 2g). In contrast to pMIC itself, the post-anesthetic period was dominated by synaptic loss (Fig. 2h), while both post-anesthetic synaptic gain and loss remained non-significantly increased. Intriguingly, a substantial fraction of synapses gained during pMIC remained stable across the 3-day post-anesthetic period (42.69 ± 10.87 % s.e.m. [total gain: 59 synapses in n=6 mice]).

**Figure 2.**
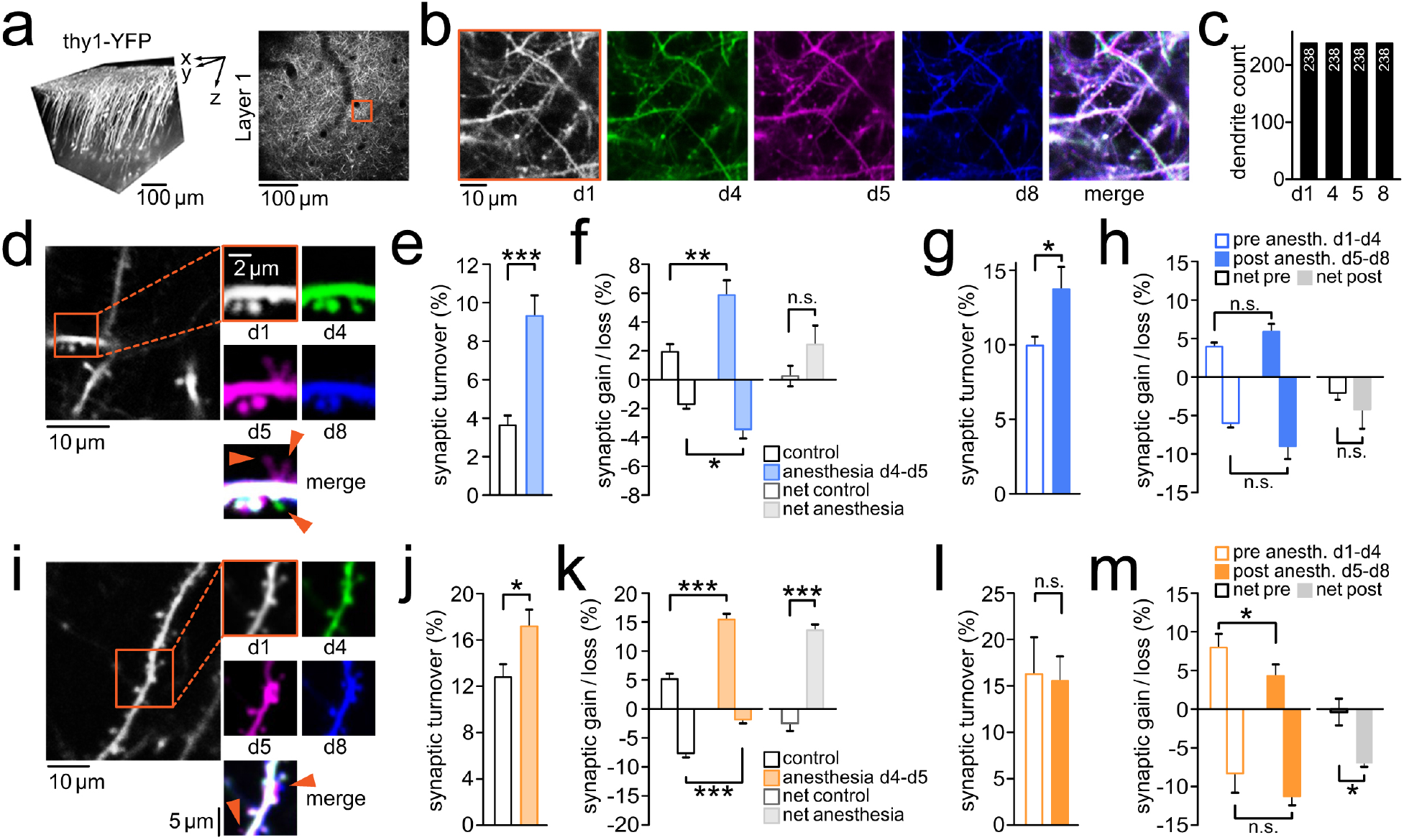
pMIC leads to synaptic alterations in adolescence and adulthood. **a)** Left: 3D reconstruction of imaged layer 5 pyramidal neurons in vivo. Imaging was typically carried out up until 150μm beneath the pial surface. Right: Axial imaging field of view (FOV) at low magnification just below the brain surface to precisely re-identify highly zoomed-in FOVs across multiple imaging sessions using vascular landmarks. **b)** red inset shown in a; identical dendritic segments were imaged 4 times before pMIC (d1, d4), and following pMIC (d5, d8) in an adult mouse, with a complete overlap of dendritic segments across imaging sessions (right). **c)** Quantification of structural dendritic changes across pMIC in adult mice (n=6); dendritic segments remained completely stable, with no dendrites gained, or lost. **d)** Paradigmatic dendritic segment with associated synaptic changes across pMIC (baseline: d1-d4, pMIC: d4-d5, post pMIC: d5-d8). Two synapses are transiently gained during pMIC (magenta), 1 synapse is permanently lost (green). **e)** Total synaptic turnover during pMIC in adult mice (n=6) vs. 1-day turnover in healthy controls (n=6); unpaired t-test: p=0.043 **f)** Synaptic gain, synaptic loss, and net-effect (gain - loss) during pMIC in adult mice (n=6) vs. healthy controls (n=6); unpaired t-test: p=0.0064 (gain), p=0.0354 (loss), p=0.1728 (net-effect). **g)** Total 3-day synaptic turnover within adult mice (n=6) during baseline (d1-d4) vs. post pMIC (d5-d8); paired t-test: p=0.0484 **h)** Synaptic gain, synaptic loss, and net-effect (gain - loss) within adult mice (n=6) during baseline (d1-d4) vs. post pMIC (d5-d8); paired t-test:p=0.4913 (gain), p=0.16 (loss), p=0.4441 (net-effect). **i)**Paradigmatic dendritic segment with associated synaptic changes across pMIC (baseline: d1-d4, pMIC: d4-d5, post pMIC: d5-d8). Two synapses are gained during pMIC (magenta), and persist during the post pMIC period (blue). **j)** Total synaptic turnover during pMIC in adolescent mice (n=4) vs. 1-day turnover in healthy controls (n=5); unpaired t-test: p=0.0433 **k)** Synaptic gain, synaptic loss, and net-effect (gain - loss) during pMIC in adolescent mice (n=4) vs. healthy controls (n=5); unpaired t-test: p=0.0002 (gain), p=0.0007 (loss), p<0.0001 (net-effect). **l)** Total 3-day synaptic turnover within adolescent mice (n=4) during baseline (d1-d4) vs. post pMIC (d5-d8); paired t-test: p=0.7099 **m)** Synaptic gain, synaptic loss, and net-effect (gain - loss) within adolescent mice (n=4) during baseline (d1-d4) vs. post pMIC (d5-d8); paired t-test: p=0.043 (gain), p=0.1671 (loss), p=0.0386 (net-effect). Depiction of statistical significance in fig. 2: *p<0.05, **p<0.01, ***p<0.001. All bar plots show mean ± s.e.m..

Next, we sought to put our findings in adult mice in developmental context by examining younger animals (~PND 30). Clinical studies, and animal research suggest that shorter MIC periods (max. 6 hrs tested to date) at young age (≤ 1month in rodents) may be associated with long-term cognitive impairment^9^, and increased synapse formation with an age-dependently decreasing effect size (no MIC effect on dendritic spines at age >1 month)^7,8,13^.

Similarly to adult mice, pMIC in young mice (n=4, pMIC duration = 18.5 ± 1.26 hrs s.e.m.) was associated with a significantly enhanced 1-day synapse turnover (Fig. 2i-j, n=4 mice, total synapse count: 284 [71 ± 12 s.e.m.]). In line with previous results on immature synapses in 1 month old mice (isoflurane for 4hrs)^13^, synaptic loss was significantly reduced during pMIC compared to controls (Fig. 2k, n=5 mice, total synapse count: 587 [117.4 ± 15 s.e.m.]). At the same time, consistent with our results for pMIC in adult mice, pMIC-associated synapse formation was strongly increased resulting in a pronounced net-positive synaptic turnover (Fig. 2k). When comparing the pre- and post-anesthetic 3-day synaptic turnover, total turnover remained unchanged (Fig. 2l). Yet, in contrast with adult mice, synaptic formation post pMIC in younger mice was significantly reduced, and synapse elimination increased, resulting in a net-negative synaptic turnover (Fig. 2m). Again, a substantial fraction of synapses gained during pMIC remained stable during the post-anesthetic period (50.57 ± 9.09 % s.e.m. [total gain: 36 synapses in n=4 mice]).

Our data demonstrate that prolonged general anesthesia alters object recognition and cortical synaptic architecture in both young and adult mice. With isoflurane, our findings point towards net-positive synapse turnover during pMIC, followed by a net-negative synapse turnover during the post-anesthetic period. Across all examined ages, a high fraction of newly formed spines during pMIC displayed unexpected post-anesthetic stability suggesting that they may form persistent synaptic connections. Given the fundamental relevance of synaptic connections and experience-driven synaptic plasticity for cognitive function and memory^19–21^, our observed alterations in pMIC-associated synaptic dynamics represent a possible factor contributing to cognitive deficits following general anesthesia. The mechanisms underlying the observed alterations remain unclear, yet likely play out across multiple anatomical scales^22^. On the molecular level, anesthetics target ion channels^23^. On the organelle level, anesthetics e.g. exert effects on a cell’s cytoskeleton^24^. Ultimately, circuit level alterations driven by anesthetics may play a main role in the pMIC-associated synaptic plasticity observed here, since the prolonged pharmacological inactivation profoundly disrupts neural circuits that under physiological conditions dynamically adapt at the synaptic level with changing intrinsic (e.g. circadian protein expression)^25^, and extrinsic environmental parameters (e.g. novel sensory experience) across age^21^.

Our findings carry profound implications for medical care, as they point out at significant structural and functional side effects of pMIC. To date, no standard approach exists to prevent cognitive side effects or alterations of brain architecture during pMIC. A better understanding of pMIC-related effects on cortical synapses, especially in the context of different widely used anesthetics, could allow for more individually tailored anesthetic regimens, and foster research on adjuvant therapeutic strategies. Such research may pave the way towards methods to improve cognitive outcomes of ICU survivors, and thus and help reduce the socioeconomic consequences of prolonged medically induced coma.

## Acknowledgments

We thank Yuste Lab members for useful comments on this project, and H.W. Pfister for clinical supervision. Supported by the NEI (R01EY011787) and NIMH (R01MH115900). This material is based upon work supported by, or in part by, the U. S. Army Research Laboratory and the U. S. Army Research Office under contract number W911NF-12-1-0594 (MURI). The authors have no competing financial interests to declare. M.W. conceived the project, and designed the experimental framework. M.W., D.S.P., and A.L. established the anesthesia setup. M.W. and A.L. performed the experiments. M.W. (clinical data/vital parameters, structural imaging, behavioral assessment), A.L. (clinical data/vital parameters, structural imaging, behavioral assessment), D.S.P. (structural imaging), and S.H. (behavioral assessment) analyzed the data. S.H. provided custom code (MATLAB, MathWorks) for automated tracking of mouse exploratory behavior. M.W. wrote the paper. All authors discussed results, and edited the paper. R.Y. assembled and directed the team and secured funding, and resources. R.Y. is an Ikerbasque Research Professor at the Donostia International Physics Center.

## Methods

### General information

All experiments were carried out in compliance with the Columbia University institutional animal care guidelines. We used B6.Cg-Tg(Thy1-YFP)HJrs/J mice^14^ (Jackson Laboratories; RRID: IMSR_JAX:003782), where cortical layer 5 pyramidal neurons are preferentially labeled. For experiments, postnatal age of mice ranged from 1 to 12 months (two age groups: p30-35, or p120-360). Mice were housed at a 12 hour light/dark cycle, and food and water was provided ad libitum.

### Surgical procedures

For repeated two-photon imaging of cortical synapses, a reinforced thin skull chronic cranial window was established as described^15^. We chose a “thin skull” approach to minimize potential craniotomy-related changes of synaptic plasticity that have been reported previously^26^. In brief, mice were anaesthesized with isoflurane (initial dose 2-3% partial pressure in air, then reduction to 1-1.5%). Prior to surgery, all mice received carprofen (s.c.), enrofloxacin (s.c.), and dexamethasone (i.m.). Throughout surgical procedures, proper anesthetic depth was checked intermittently by toe, and tail pinching. Under sterile conditions, a small flap of skin above the skull was removed and a titanium head plate with a central foramen (7×7mm) was fixed on the skull using dental cement. Then, a dental microdrill was used at low rotation speed to carefully thin a small region (usually <1mm in diameter) over the left somatosensory cortex. To prevent the skull and brain tissue from heating up by the drilling process, drilling and wetting with sterile artificial cereberospinal fluid (ACSF) were alternated frequently. At a thickness of around 50μm, the skull typically began to bend upon gentle pressure and the intracranial vasculature became clearly visible under the moistened skull. In addition, small air bubbles within the thin spongioform part of the thinned skull appeared when ACSF was applied. From here on, the skull was further thinned at an even lower drilling rotation speed and with minimal vertical pressure, down to a thickness of 10-20 μm. When thin enough, the small air enclosures in the skull no longer appeared upon ACSF application.

At this point, the skull was allowed to dry completely. Once dry, a small drop of cyanoacrylate glue was applied onto the thinned skull. Immediately following the application of the glue, a thin glass cover slip (3mm in diameter, No. 0, Warner Instruments) was lowered, and gently pressed onto the thinned area, held in place by a stereotactic arm. After glue solidification (several minutes), the edges of the glass cover slip were sealed off by dental cement. Following the surgical procedure, mice received pain medication, and were allowed to recover for at least 3 days. The “reinforcement” of the cranial window prevented the skull from growing back making it possible to image the same cortical structures repeatedly without the need of re-thinning the skull.

### Experimental timeline

The experimental timeline is displayed in figure 1 d. Following recovery from surgical establishment of a chronic cranial window, mice were acclimatized to the experimenter, until no signs of distress were present (usually ~3 sessions, 30min each). Then, two imaging sessions were carried out to determine the baseline 3 day (‘d’) synaptic turnover (d1, d4). In between baseline imaging sessions, mice were habituated to the open field maze (d3), and subjected to a first novel object recognition test^16^ (NOR1, d4). Immediately post imaging on day 4, pMIC was initiated. To document synaptic changes occurring during pMIC, a third imaging session was carried out right at the end of pMIC (d5). After at least 24 hours of recovery from pMIC, mice were subjected to a second NOR (NOR2, d7). Three days after pMIC termination, mice were imaged once again to determine their post-anesthetic 3 day synaptic turnover (d8).

### Prolonged medically induced coma

The tightly controlled and safe pMIC procedure (−24hrs) required the continuous presence of an experimenter. Mimicking hospital conditions, remote video and vital parameter monitoring allowed the experimenter to leave the pMIC setup while maintaining the ability to rapidly carry out setup adjustments. Based on a protocol of long-term anesthesia in mice^27^, we established a setup that permits continuous general anesthesia of mice for up to days (Fig. 1a). While anesthetized, mice were kept on a warming pad maintaining a body core temperature of 37.5°C through a rectal probe and closed-loop temperature control system (ATC2000, World Precision Instruments). The warming pad was placed on a tiltable platform whose angle was slightly changed intermittently (max. 4°) in order to shift the animals center of mass to reduce pressure pain and bruising. As mice loose body temperature easily due to their anatomy, an infrared lamp was additionally put in place and optionally switched on, if needed. Isoflurane was delivered in humidified air through a custom nose piece (initial dose 1.5-2.5% partial pressure in air [ppa], steady state dose 0.5-1.5% ppa). Ventilation air was humidified between the isoflurane vaporizer and the nose piece, and all waste gas was actively removed via a suction pump. By the help of a dual gas flowmeter system (100% O_2_, and room air) set up before the isoflurane vaporizer system, the oxygen fraction (FiO_2_) in air could be modified seamlessly between 21-100%. FiO_2_ adjustment depended on the measured peripheral arterial O_2_ saturation. During the early period of pMIC, a fraction of inspired oxygenation (FiO_2_) of 21% was completely sufficient to maintain high peripheral O_2_ saturation in all anesthetized mice, while subsequently (>8 hrs) FiO_2_ was usually slowly increased towards pMIC termination without exceeding 50%. Similarly to human intensive care, we tried to avoid an fiO_2_ >50%^18^. To provide the animal with sufficient fluid and nutrition during anesthesia, continuous fluid (saline) and nutritional supply (1% glucose solution, in PBS) was administered via a programmable perfusion system via a subcutaneous line at a rate of 0.1-0.2ml/hr. All throughout anesthesia, basic vital parameters (blood oxygenation, heart rate, breath rate, pulse distension, breath distension) were continuously monitored and recorded by a thigh or paw sensor (mouse ox plus system, Starr life sciences). The sensor was periodically moved from one extremity to another to avoid pressure-induced tissue damage. Further, this approach minimized the risk of faulty vital parameter sensing due to excessive tissue compression. A sustained pulse/breath distension ratio below 1 was sought to be avoided^27^, as it indicates anesthesia being too deep (large inhalatory excursions, gasping; in combination with brady- or normocardia), or a lack of intravascular fluid (hypovolemia; in combination with tachycardia). The experimenter was continually present in the laboratory to perform imaging immediately prior and post pMIC, and to adjust for example anesthetic depth or fluid supply with minimal delay all throughout anesthesia. Real-time streaming of video monitoring (Thorlabs, DCC1645C) and vital parameters through a local network allowed for remote monitoring of anesthetized animals.

### Two-photon imaging

For chronic surveillance of structural synaptic changes in vivo, dendritic arborization, and dendritic spines of layer 5 cortical neurons expressing yellow fluorescent protein (YFP)^14^ were imaged 4 times (day 1, day 4 [start of anesthesia], day 5 [end of anesthesia], day 8) using a two-photon microscope (Bruker Ultima; Billerica, MA) and a Ti:Sapphire laser (Chameleon Ultra II; Coherent) at 940 nm through a 25x objective (water immersion, N.A. 1,05, Olympus). Galvanometer scanning and image acquisition was controlled by Prairie View Imaging software. Each imaging session (~15-30 minutes) included 1-4 high resolution z-stacks in different locations (field of view typically ~50 x 50 μm, 512 x 512 pixels, step size 1.5 μm, stack depth up to 150 μm beneath the pial surface). Imaged z-stacks were precisely co-registered across each imaging session by vascular landmarks and several different optical zoom average images of the cortex. For each imaging session, head-restrained animals were kept under light isoflurane anesthesia (0.5-1% partial pressure in air) via a nose piece while body temperature was maintained with a warming pad (37.5°C).

### Novel object recognition test

Before and after anesthesia, mice were subjected to cognitive testing by use of a novel object recognition test (NOR), a widely used basic measure of cognitive function and recognition memory in mice^16^. Following habituation (exp. day 3, figure 1 d) to an open field maze (50×50 cm), mice underwent a first NOR (exp. day 4, prior to start of pMIC). Initially, mice were exposed to two identical objects (e.g. cell culture flasks filled with sand, or LEGO towers) that were placed in the open field for 10 minutes (“familiarization”). After familiarization, mice returned to their home cage for 4-5 hours, upon which they were placed in the open field maze again, yet with one of the two identical objects replaced by a novel object (cell culture flask, or lego tower, respectively). The NOR was repeated with a new set of objects at least 24 hours after pMIC termination (exp. day 7), so changes in basic cognitive function, specifically recognition memory, could be evaluated. All behavioral testing was video-recorded (7.5 fr/sec) from above the open field. We used custom written Matlab code for automated tracking (travelled distance, object exploration time) of mouse exploratory behavior during NOR (Fig. 1c).

### Data analysis

Similarly to previously described procedures for quantifying synapse dynamics^28^, stacks were analyzed using the freely available software ImageJ (rsbweb.nih.gov/ij/). Individual 3d-stacks for each time point (d1, d4, d5, d8), and imaged areas (1-4 per animal) were analyzed side-by-side by two examiners who were blinded to conditions, except for the reference imaging timepoint (d1). Based on consensus between the two blinded examiners, clearly visible protrusions on dendritic segments were classified as synapses. For each dendritic segment analyzed, same synapses were defined as present (1) or absent (0) across stacks (imaging time points). Based on the examiners’ consensus, synapses could be defined as stable, gained, or eliminated across imaging time points, once an entire experiment was analyzed, and imaging time points could be revealed, and thus re-assigned to individual stacks. Statistical differences between synaptic turnover, synaptic gain, or loss were tested using paired (pre-vs. post-anesthetic), or unpaired (control group vs. anesthetized group) student’s t-test. All statistical analyses were carried out using MATLAB-R2017b (MathWorks), and Prism 5 (GraphPad Software).

### Data availability statement

The data that support the findings of this study are available from the corresponding author upon reasonable request.

### Code availability statement

The custom MATLAB code for automated behavioral tracking is available at: https://github.com/hanshuting/mnovobj

## Notes

**Conflict of Interest:** The authors declare no competing financial interests

